# AI4CellFate: Interpretable Early Cell Fate Prediction with Generative AI

**DOI:** 10.1101/2025.05.12.653464

**Authors:** Inês Cunha, Luca Panconi, Sebastian Bauer, Maxime Gestin, Emma Latron, Erik Sahai, Alix Le Marois, Juliette Griffié

## Abstract

Live-cell imaging provides a unique insight into complex cellular processes including single cell fate, but remains limited by both low-throughput and the lack of generalisable analytics for the multidimensional datasets it produces. This work introduces AI4CellFate, an interpretable and data-driven machine learning framework for predicting cell fate from microscopy timelapses, applied here to cancer therapy. By integrating generative AI and contrastive learning, AI4CellFate enables early fate prediction as well as visualisation of biologically relevant features, with limited annotation.

## Main Text

Live-cell microscopy imaging has proven to be an essential tool to understand complex cellular processes including division, migration and differentiation. As such, it has become instrumental for investigating drug responses *in vitro*, including cancer treatment^1^. However, live-cell imaging is low-throughput, requires costly experimental setups, and leaves cells susceptible to phototoxicity and photobleaching^2^ (Figure 1b). Following acquisition, existing analysis strategies rely on hand-picked and hypothesis driven feature extraction^3^, which introduce human bias and lead to information loss. Additionally, extensive manual annotation is often required, making the analysis process labour-intensive^4^ (Figure 1c).

**Figure 1.**
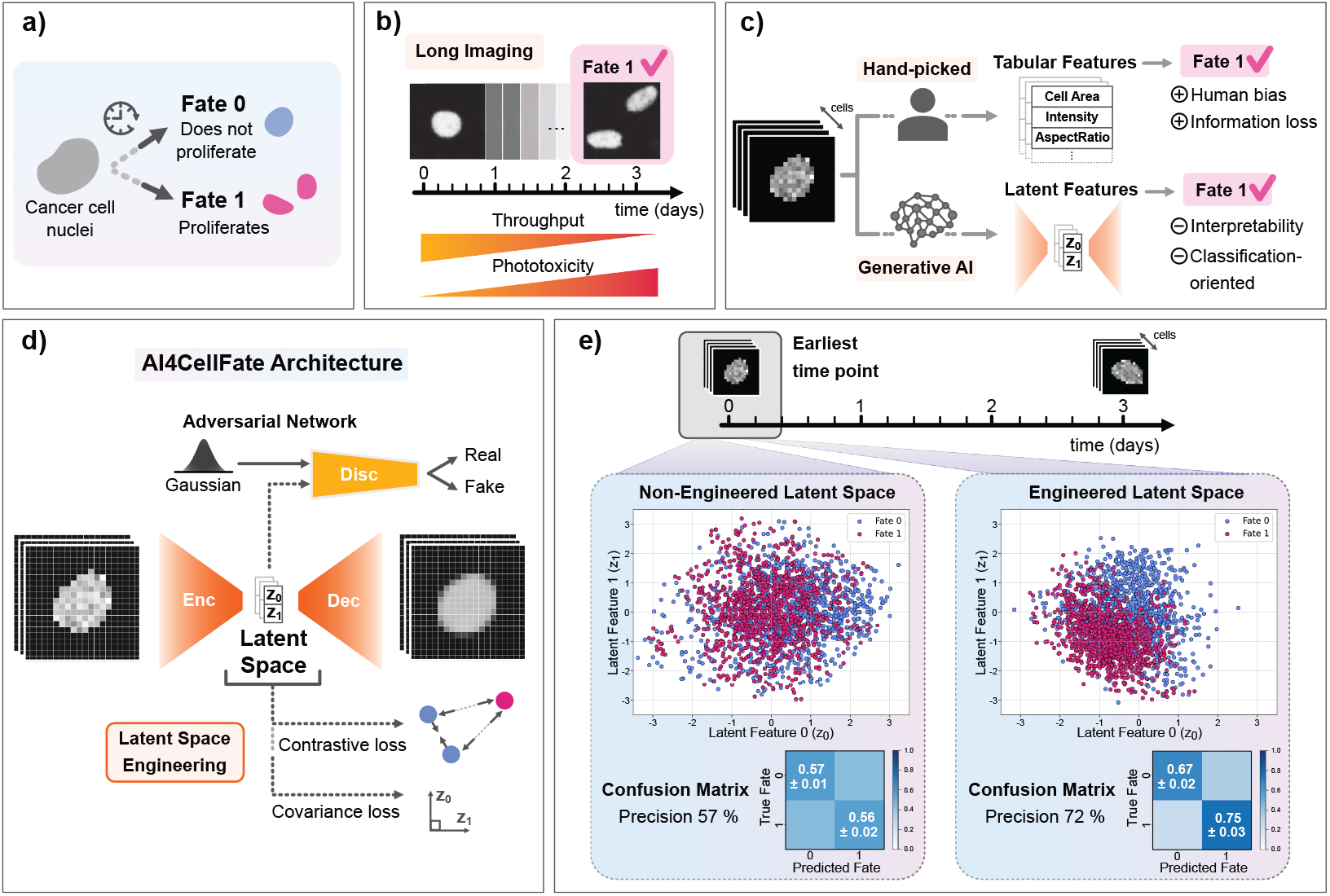
Limitations in studying live-cells and how AI4CellFate addresses them. a) Schematic of cell fates. b) Schematic of experimental limitations associated to long imaging. c) Schematic of analysis limitations associated to cell fate prediction. d) AI4CellFate architecture. e) AI4CellFate performance with and without LS engineering on the earliest frame.

Recently, generative AI has been successfully used for data-driven exploration. Autoencoders^5^, a type of generative AI, reduce input image data into a lower-dimensional space - the latent space (LS) - providing an alternative to hand-picked feature extraction. Researchers have leveraged LS representations to study images^6–9^, and recent advances suggest that the LS can be fine-tuned for human interpretability^10–12^. Whilst promising, if *in vitro* single cell drug response assays are to become a routinely used tool to generate biomedical novelty, these require dedicated analytics for high throughput and interpretable data-driven early prediction.

Here, we propose AI4CellFate, an interpretable generative AI method optimised for prediction of cell fate from live-cell imaging. We applied this model to investigate cancer cell responses to therapy, with a focus on cell phenotypes evading treatment, also called persister cells. Our dataset includes lung cancer cells exposed to a growth inhibition treatment, labelled with a nuclear-localised FRET biosensor to measure the activity of a kinase involved in proliferation and survival^3^ (see Supplementary Note 1). In response to this treatment, some cells continue to proliferate (fate 1), while others do not (fate Figure 1a).

AI4CellFate learns interpretable features from single-cell images, while ensuring they are optimised for fate prediction. The model consists of an autoencoder architecture, including an encoder, which compresses images into a latent space, and a decoder, which aims to reconstruct the original image using only latent features. This approach bypasses human bias by forcing the model to learn an efficient representation of the data in which only the most important features of the input images are retained without any *a priori* knowledge. In particular, we used an adversarial autoencoder (AAE)^13^, which uses a discriminator to generate the latent space (Figure 1d) (see Methods and Supplementary Tables 1-4). The latent space produced by an AAE has been shown to achieve better performance when used for classification tasks than the alternative variational autoencoder architecture^13^. Here, the classification task involves predicting cell fate from single frames of the live-cell movie. To increase predictive accuracy whilst enabling model interpretability, we engineered the latent space via a multi-objective loss function. This consisted of the weighted sum of four losses, which were optimised simultaneously: reconstruction loss – from the autoencoder; adversarial loss – from the discriminator; covariance loss^14^ – which ensures latent feature independence; and contrastive loss^15^ – which enhances class separation in the latent space (Figure 1d). Specifically, we used supervised contrastive learning^15^, where samples from the same class (cells with the same fate) are pulled closer together in the latent space, while those from different classes are pushed apart (see Methods). Here, each image was represented using only two latent features (z_0_ and z_1_), as this gave the best compromise for both classification and interpretable latent representation (see Supplementary Note 2, Supplementary Figure 1). The model comprised <10^6^ parameters, enabling fast training for most standard laptops.

We applied AI4CellFate to cell nuclei segmented from single-frame microscopy images extracted from live-cell timelapses. Each cell is captured within a 20×20 pixel (i.e., ∼25×25µm^2^) image and a total of 2184 cells were used (see Methods). Initially, we trained AI4CellFate using the earliest time point of the experiment. In Figure 1e, we show the obtained LS representations with and without LS engineering. Here, each cell is represented by a point in the LS. In the context of prediction, we observe increased separation between the fates in the LS when covariance and contrastive losses are added (Figure 1e, right and Supplementary Figure 2). Consequently, there is a substantial increase in model accuracy and precision (see Methods) for predicting cell fate (57% to 72%). Strikingly, our results demonstrate that cell fate could be retrieved from the first frame. For the purpose of interpretation, we confirm that the latent features are uncorrelated (see Supplementary Figure 1), and the LS follows a Gaussian distribution (see Methods). In addition, we compared AI4CellFate to the state-of-the-art approaches that use hand-picked features from the segmented cells, also on the first time point (see Methods, Supplementary Note 3, Supplementary Figure 3), showing we systematically outperform these. Moreover, hand-picked features led to significantly varied performance depending on the chosen pair of features, highlighting the impact of human bias and reproducibility issues.

We then investigated the impact of the chosen single-frame (i.e. choice of time point) on model performance. Model precision and accuracy were recorded across the normalised cell relative time - that is, from the first frame of the acquisition until death or mitosis for each cell (Figure 2a). Results suggest that both precision and accuracy remain stable across the entire cell lifetime (between 70 and 80%).

**Figure 2.**
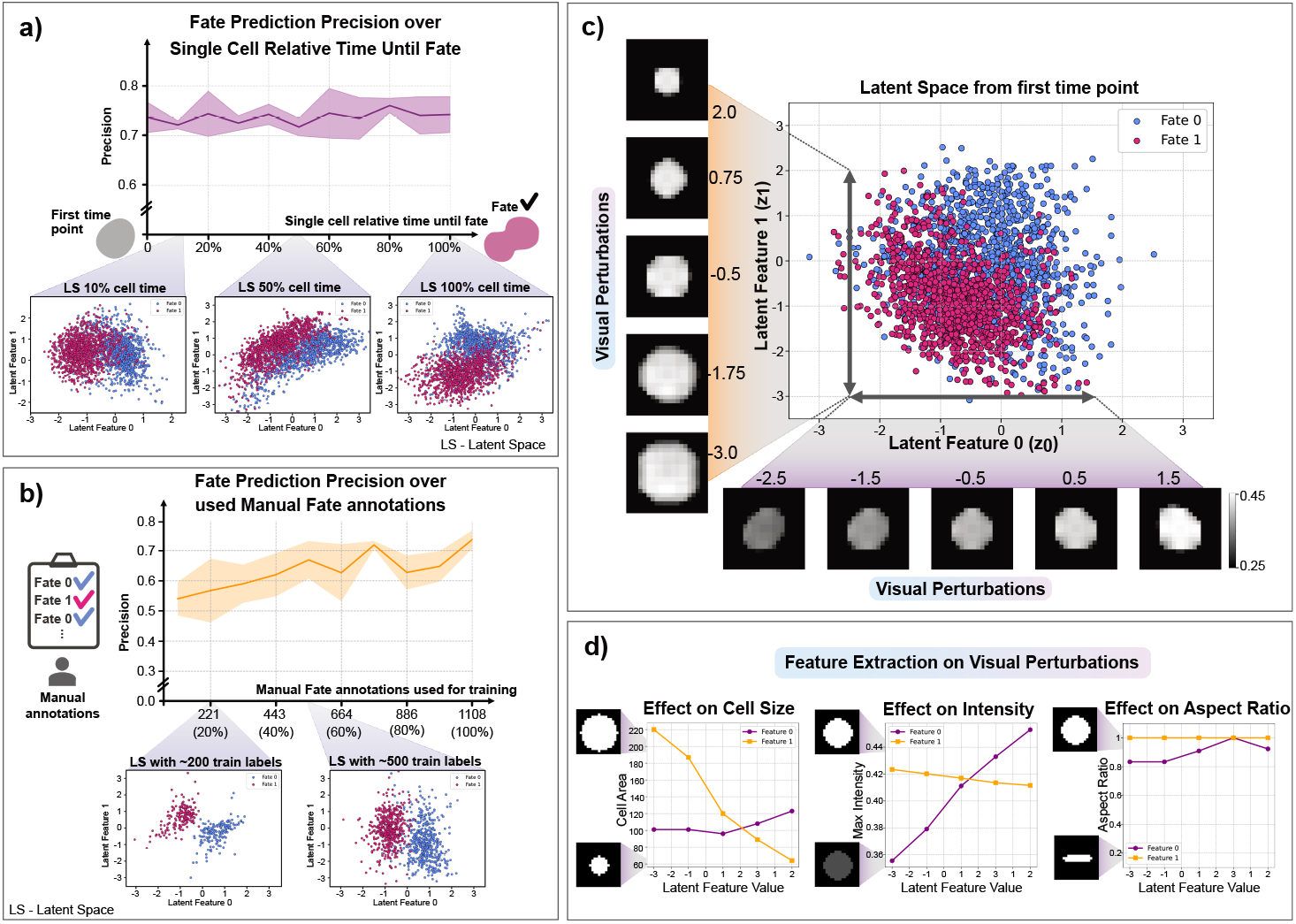
AI4CellFate Performance and Latent Feature Interpretation. a) Model precision along normalized single cell relative time until fate. b) AI4CellFate performance over number of annotated data for training. c) Visualisation of LS perturbations along dimensions (generated images). d) Feature extraction from the perturbations.

Next, we assessed how the amount of annotated data in the training set affected model performance, since AI4CellFate enables on-the-fly visualisation of the class separation in the LS. We found that, on the applied unsynchronised cancer cells, which are associated with highly complex biological variability, AI4CellFate can already learn meaningful representations, achieving class separation and generalisation with as little as 500 annotated cells (Figure 2b).

To interpret the biological significance of learned latent features, we performed perturbations along each LS dimension by fixing one feature and varying the other (Figure 2c). Using the AI4CellFate decoder, we were able to visually interpret the impact of each feature in generated nuclei images. We observed that feature z_0_ correlates with signal intensity, while feature z_1_ correlates with cell size. These findings are validated by performing feature extraction on perturbed images (Figure 2d). Biologically, this translates to fate 1 (i.e., proliferating cells) being associated with higher ERK activity measured by FRET (see Methods), and greater overall cell size, as supported by literature^3^. Notably, these findings were concluded from the earliest time point of the experiment.

In summary, AI4CellFate provides a fully data-driven framework for predicting cell fate from single frames of time-lapses. By leveraging multi-objective loss function for LS optimisation, AI4CellFate enables visualisation and interpretation of the cellular features affecting fate predictions (i.e., why could the model predict reliably). Here, we applied AI4CellFate to cancer treatment response, identifying key features for the prediction of cell fate in an unbiased manner. Using a single, early time point (i.e., frame) as input data provided similar predictive power to later time points, suggesting that persister cells commit to division a long time before mitosis actually occurs, and that the signalling and morphological traits associated with future division are stable throughout the cell cycle. We demonstrate that from the earliest experimental time point and with few manual annotations, we can extract reliable and relevant biological insights.

AI4CellFate enables early cell fate prediction, which leads to higher throughput and can easily be integrated in a smart microscopy framework^16^. Furthermore, by enabling the visualisation of features that contributed to prediction, AI4CellFate provides an unbiased platform for interpretability. This paves the way for novel insights into cellular processes such as drug response, disease progression, or treatment resistance. More broadly, this approach moves beyond hypothesis-driven frameworks that typically focus on preselected single markers, embracing instead data-driven methodologies (i.e., no *a priori* knowledge) that leverage high-content, multi-dimensional image acquisition. We anticipate that approaches such as AI4CellFate will provide a catalyst for novel discoveries.

## Methods

### Image preprocessing

The dataset used in this study consists of live-cell images of lung cancer cells labelled with a FRET biosensor that measures ERK/MAPK activity in the cell nuclei, collected across three biological replicates^3^. Further details on the dataset can be found in Supplementary Note 1. From the raw images, multiple cellular features were previously extracted for each cell at each time point^3^. These features included cell position, the average and standard deviation of CFP and YFP (channels 1 and 2) intensities, cell-wide FRET intensity and intensity sum, as well as morphological properties such as major and minor axis length, eccentricity, diameter, perimeter, area, and aspect ratio. Additionally, cell segmentations were provided, along with manually annotated cell fate and mitosis time as extra features. In total, the dataset comprised 1,827 cells, tracked over a maximum of 1,080 time points, with 19 extracted features in tabular format.

To prepare the data for model training, several preprocessing steps were applied. First, a 20×20 pixel sub-image centred on each cell nucleus was extracted for all time points. Cells that did not survive for the entire 1,080 time points (either undergoing mitosis or cell death) were padded with zeros for the remaining frames.

To remove background noise, all images were overlaid with their corresponding segmentation masks. To prevent bias in model decision-making, only first-generation cells were retained. Daughter cells were removed by filtering out any cells that were not present at the start of imaging and by excluding post-mitosis frames within the same mother cell track. Additionally, cells located at the edges of the field of view and those in close proximity to other cells (i.e., where segmentation masks merged multiple nuclei) were excluded.

Following preprocessing, the final dataset consisted of 1,385 cells, each tracked over 1,080 time points, with 2 imaging channels (CFP and YFP) and an image height and width of 20×20 pixels. The corresponding tabular dataset contained the same 1,385 cells with 19 extracted features. However, for model training on tabular data, only 13 features were retained, excluding cell ID, x/y position, time point, fate, and mitosis time.

For image data, the first channel (CFP) was divided by the second channel (YFP), generating “FRET-like” single-channel images representing the inverse of ERK/MAPK activity. These images were then normalised by dividing by the dataset-wide maximum intensity to maintain variability across cells. Additionally, pixel intensities were stretched (to a minimum value close to zero) to enhance variations while preserving the relative intensity distribution within and between cells.

#### Data Splitting and Augmentation

To ensure a balanced representation of cell fate classes in both the image and tabular data, we performed a stratified train-test split. The dataset was split into 80% training and 20% testing using stratified sampling based on cell fate labels to maintain class proportions. This yielded 1108 cells in the training set and 277 cells in the test set. The split was performed using a random state of 42.

The dataset exhibited class imbalance, with only 17% of cells belonging to fate 1 (proliferating cells). To mitigate this imbalance and improve model performance, data augmentation was applied to the training image dataset. Augmentations were performed on both fate 0 and fate 1 cells, generating multiple transformed versions of each time-series movie to increase variability while preserving biological relevance. Each movie underwent random horizontal and vertical flips, as well as 90°, 180°, and 270° rotations to ensure that orientation did not bias the model. To balance the dataset, cells with fate 0 were randomly selected to match the same number of cells as fate 1. Finally, the dataset was shuffled to eliminate any ordering biases before training. This yielded a training image set of 2184 cells (equally distributed in fate 0 and 1).

#### Single-cell relative time normalisation

To study cell behaviour at different stages of their lifetime, we applied single-cell relative time normalisation, which standardised time points across all cells. Instead of analysing cells at fixed absolute time points, we normalised each cell’s time progression relative to its cell fate, starting from the first acquired frame. For the tabular data, we identified each cell’s recorded lifetime i.e., starting with the first acquired frame, by determining the number of nonzero values in a given feature, as these corresponded to the number of time points for which the cell was alive/ non-splitting for (since the rest was padded with zeros). Feature values were then extracted at 11 evenly spaced time points (0%, 10%, …, 100%) of each cell’s lifetime. For the image data, we applied a similar normalisation strategy. We determined each cell’s recorded lifetime based on the number of nonzero image frames (frames that contained nonzero pixel values). Then, we selected images at 11 normalised time points. This allowed us to compare cells at equivalent life stages, despite differences in absolute time duration.

### Model architecture

The adversarial autoencoder (AAE) employed in this study consists of three main components: an encoder, a decoder or generator, and a discriminator. The detailed architecture of each component is provided in Supplementary Tables 1-4. The AAE is designed to learn a latent space representation of input images. This is a 2-dimensional latent space, i.e., with 2 features representing each input image. More information about the chosen latent space dimensionality can be read in Supplementary Note 2 and Supplementary Figure 1.

The encoder maps input images into the low-dimensional latent space, using two residual downsampling blocks with a series of convolution layers with spectral normalisation to ensure stability during training^17^. The residual block also applies Gaussian noise layers (standard deviation = 0.003) to enhance generalisability of the learned features.

The decoder uses the produced low-dimensional latent space from the encoder as input to reconstruct images. The latent representation is progressively upsampled with residual upsampling blocks. The output is a reconstructed image with the same dimensions as the input.

The discriminator serves as an adversarial component, distinguishing between the true latent representations and “fake” generated ones. It consists of a feedforward neural network composed of three fully connected layers. The last layer outputs a single neuron with a sigmoid activation, which predicts whether a given latent vector originates from the true encoding distribution or is artificially generated.

The learned latent space is subsequently used for classification with the classifier detailed in Supplementary Table 5, which consists of a single dense layer with two output neurons for classification into the two cell fates.

### Training

The training procedure optimises a multi-objective loss function, which is the sum of four loss terms – reconstruction, adversarial, contrastive and covariance loss-weighted by their respective lambda coefficients.

#### Reconstruction Loss

The first loss is the reconstruction loss, which is a multi-scale structural similarity (MS-SSIM)^18^ loss between the original and reconstructed images. MS-SSIM captures perceptual image quality better than pixel-wise losses like mean squared error, as it considers structural information at multiple scales. This makes it more robust to small variations in intensity^19^. It is computed as a weighted sum of SSIM values across multiple scales, as follows. For two images x and 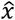, the SSIM at a single scale is computed as:

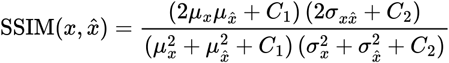

where 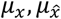 are the local mean intensities of x and 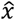, computed using a gaussian filter with a kernel size of 5 and a sigma of 1. 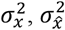 are the local variances and 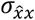 is the local covariance. *C*_1_ and *C*_2_ are small constants for numerical stability, calculated based on the pixel intensity range. MS-SSIM then extends SSIM by computing it at multiple scales. The images are progressively downsampled, and SSIM is computed at each scale:

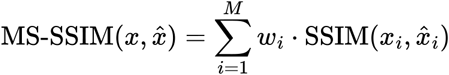

Where M=5 is the number of scales and *ω*_*i*_ are the scale dependent weights of [0.0448, 0.2856, 0.3001, 0.2363, 0.1333].

The final reconstruction loss function is defined as:

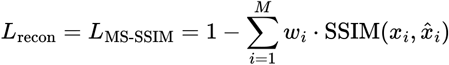

This ensures that maximising the similarity score minimises the loss.

#### Adversarial Loss

The second loss is an adversarial loss from the discriminator, consisting of a binary cross-entropy (BCE) loss for the discrimination between real latent samples and “fake” generated Gaussian samples. This ensures that the latent space aligns with a Gaussian distribution, enforcing a structured representation useful for downstream tasks.

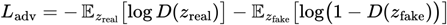

Where *z*_*real*_ are latent representations from real images and *z*_*fake*_ are sampled from a prior Gaussian distribution, and *D*(*z*) is the discriminator that predicts whether *z* is real or fake.

#### Contrastive Loss

The third loss is the contrastive loss, which encourages class separation in the latent space by bringing similar samples closer together in distance, while pushing dissimilar samples apart. In the context of supervised contrastive learning^15^, this helps in learning a discriminative latent space that retains important information for the classification task. In this case, it is used to ensure that samples with the same fate are closer together in the latent space while maintaining sufficient separation from the other fate.

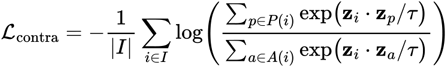

Where *z*_*i*_ is the normalised latent representation of sample (anchor) *i, τ* is the temperature value (set to 0.5), *P*(*i*) is the set of positive samples (i.e., samples from the same fate as the anchor), and *A*(*i*) is the set of all samples except the anchor (full batch minus *i*). This means that the numerator sums over all positive pairs’ similarities, and the denominator sums over all samples’ similarities.

#### Covariance Loss

The fourth loss is the covariance loss, which forces the latent features to be independent by minimising off-diagonal elements of the covariance matrix of the latent representation^10,14^. This regularisation prevents redundancy among latent dimensions, ensuring that each latent feature encodes distinct and meaningful information about the images.

These losses are all sum together into the total loss function, combining all the individual losses with their respective weights (*λ*):

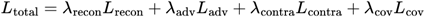

We used *λ*_*recon*_ = 8, *λ*_*adv*_ = 4, *λ*_*contra*_ = 6 and *λ*_*cov*_ = 0.0001. The weights were determined empirically, where each *λ* was adjusted based on its impact on the model’s performance. Higher weights were assigned to the contrastive loss and reconstruction loss, as they had the most significant influence on feature separation and representative reconstructions. The covariance loss was given a lower weight, as it was sufficient to ensure that the latent features remained uncorrelated without introducing instability into the training process.

#### Training procedure

Each training epoch consists of multiple mini-batch updates. In the same epoch, the autoencoder is trained first, and the discriminator is separately updated. The training steps in each epoch consist of:

1. The encoder maps the input images to a latent space *z*_*img*_
2. The decoder takes as input *z*_*img*_ and reconstructs them back into images
3. All losses are then computed as detailed above, where the reconstruction loss uses the original and reconstructed images; for the adversarial loss, the discriminator is used to classify the *z*_*img*_ as real or fake; the contrastive loss is calculated given the *z*_*img*_ and the true fate labels of each batch; and the covariance loss is calculated from *z*_*img*_. The losses are then all summed together with the corresponding weights *λ* (multi-objective optimisation).
4. The model gradients are computed and the autoencoder’s parameters are updated using backpropagation.
5. In parallel with training the autoencoder, the discriminator is trained to distinguish between real and fake latent vectors. Random vectors from a normal distribution are generated and passed through the discriminator to compute the discriminator’s loss, which is the average of the BCE loss on real and fake samples.
6. The discriminator weights are updated through backpropagation.

An Adam optimizer was used for training the encoder, decoder, and discriminator. The batch size was set to 30, and the learning rate was set to 0.001. For reproducibility, the random seeds were set to 42, 43, and 44, with the results presented in the main figures corresponding to seed 42. The number of training epochs was determined using a stopping condition based on specific metrics: (1) latent space class separation (≥1 pixel Euclidean distance), (2) KL divergence in both latent dimensions (see next section), indicating a near-Gaussian distribution (≤0.2), and (3) model accuracy and precision (≥0.65). Training stopped once all criteria were met, and the number of epochs was over 50. If the criteria aren’t met, the model stops running at 100 epochs. For seed 42, 60 epochs were enough for all stop criteria to be met. The AI4CellFate model was trained in two stages. First, the adversarial autoencoder was trained for 15 epochs with contrastive and covariance losses disabled (λ values set to zero) and *λ*_*recon*_ set five times higher than *λ*_*adv*_. This ensures stable reconstructions (reconstruction loss convergence) before introducing additional constraints to the latent space (i.e., contrastive and covariance losses). In the second stage, all losses were activated with the previously defined λ values, and training continues until the stopping criteria are met. Loss plots for both stages are shown in Supplementary Figure 4.

### Latent Space Evaluation

After training, the latent space was evaluated to ensure it follows a Gaussian distribution by computing the Kullback-Leibler (KL) divergence for both dimensions. KL divergence measures how much one probability distribution deviates from another, with lower values indicating closer alignment. In this case, it quantifies the difference between the learned latent space distribution and a normal Gaussian. Since KL divergence was part of the stopping criteria, all models achieved values below 0.2. For seed 42 at 60 epochs, the KL divergence was 0.19 for dimension 0 and 0.13 for dimension 1.

The correlation between latent features was quantified to verify the effectiveness of the covariance loss, yielding a correlation of 0.06 between the two latent features. This confirms that the loss function successfully minimized feature dependence (Supplementary Figure 1, panel b).

To assess the AI4CellFate model’s accuracy in predicting cell fates, a test latent space was generated using the trained encoder, and a confusion matrix was computed. Accuracy ((TP + TN) / 2) and precision (TP / (TP + FP)) were used as evaluation metrics. Precision was particularly emphasised for reasons detailed in Supplementary Note 4.

### Visual Interpretability

To explore the interpretability of the AI4CellFate model, we performed controlled perturbations in the latent space and analysed their effects on the reconstructed images. This approach allowed us to assess how variations in individual latent features influenced the generated synthetic cells, which translate to cellular characteristics.

We began with a baseline latent vector initialised to zeros (2,2), representing a neutral state. To examine the impact of each feature, we systematically varied one latent dimension while keeping the other constant. Specifically, perturbations were applied to feature 0 using values from [-2.5, 1.5] and to feature 1 using values from [-3, 2], each sampled at five evenly spaced points.

For each perturbed latent vector, the trained decoder generated a corresponding synthetic image. The visual perturbations are shown in Figure 2c.

To quantitatively assess these changes, key morphological features were extracted from the generated images. Specifically:

- Cell area: Defined as the number of pixels above a set intensity threshold of 0.1.
- Maximum intensity: The highest pixel intensity in the image.
- Shape descriptors: Including aspect ratio (width-to-height ratio of the bounding box) and circularity (measure of roundness based on area and perimeter).

### Tabular Data Prediction

The predictive power of hand-picked tabular features and how these compare with AI4CellFate was studied, in the earliest experimental time point (Supplementary Figure 3). A total of 13 features were extracted directly from the cell images (as explained in the image preprocessing section). A total of 78 2-feature combinations (see Supplementary Table 7 for all feature combinations) and each feature combination was evaluated with two classifiers: 1) a more “complex” multi-layer-perceptron (see Supplementary Table 6) optimised for this data and 2) the simplest multi-layer-perceptron, matching the one used for the latent features from AI4CellFate. Both precision and accuracy were calculated for each feature combination. The optimised MLP was also used on all 13 features and the same metrics were compared with the ones obtained from AI4CellFate. Discussion on the obtained results can be seen in Supplementary Note 3.

## Code Availability

All code developed in this manuscript can be accessed in the GitHub repository https://github.com/ComputationalMicroscopy4CellBio/AI4CellFate

## Author Contributions

J.G. supervised the project. I.C., A.L.M and J.G. designed the project. I.C. implemented the project. A.L.M. acquired the experimental data. I.C. and L.P. tested and optimised the code. I.C. performed validation. I.C., L.P. and J.G. wrote the manuscript. I.C., L.P., S.B., M.G, E.L., E.S., A.L.M and J.G. revised the manuscript.

## Supplementary Tables

**Supplementary Table 1:**
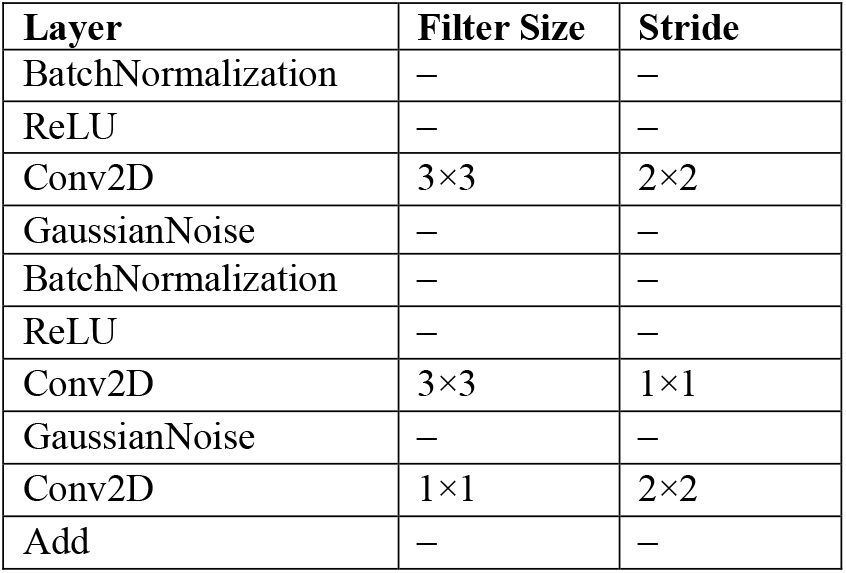
Residual Block Architecture

| Layer | Filter Size | Stride |
| --- | --- | --- |
| BatchNormalization | — | — |
| ReLU | — | — |
| Conv2D | 3×3 | 2×2 |
| GaussianNoise | — | — |
| BatchNormalization | — | — |
| ReLU | — | — |
| Conv2D | 3×3 | 1×1 |
| GaussianNoise | — | — |
| Conv2D | 1×1 | 2×2 |
| Add | — | — |

**Supplementary Table 2:** Encoder Architecture

| Layer | Input Shape | Output Shape | Filter Size | Stride | #Params |
| --- | --- | --- | --- | --- | --- |
| Input | (20, 20, 1) | (20, 20, 1) | — | — | 0 |
| Conv2D | (20, 20, 1) | (20, 20, 16) | 3×3 | 1×1 | 176 |
| ResidualBlockDown | (20, 20, 16) | (10, 10, 32) | — | — | 14720 |
| Dropout (0.3) | (10, 10, 32) | (10, 10, 32) | — | — | 0 |
| ResidualBlockDown | (10, 10, 32) | (5, 5, 64) | — | — | 58112 |
| Dropout (0.3) | (5, 5, 64) | (5, 5, 64) | — | — | 0 |
| Flatten | (5, 5, 64) | (1600) | — | — | 0 |
| GaussianNoise | (1600) | (1600) | — | — | 0 |
| Swish | (1600) | (1600) | — | — | 0 |
| Dense | (1600) | (2) | — | — | 3204 |
| GaussianNoise | (2) | (2) | — | — | 0 |
| BatchNormalization | (2) | (2) | — | — | 8 |
| Total Trainable Parameters: 75622 |  |  |  |  |  |

**Supplementary Table 3:** Decoder Architecture

| Layer | Input Shape | Output Shape | Filter Size | Stride | #Params |
| --- | --- | --- | --- | --- | --- |
| Input | (2) | (2) | — | — | 0 |
| Dense | (2) | (1600) | — | — | 6400 |
| GaussianNoise | (1600) | (1600) | — | — | 0 |
| Reshape | (1600) | (5, 5, 64) | — | — | 0 |
| ResidualBlockUp | (5, 5, 64) | (10, 10, 64) | — | — | 78720 |
| Dropout (0.3) | (10, 10, 64) | (10, 10, 64) | — | — | 0 |
| ResidualBlockUp | (10, 10, 64) | (20, 20, 32) | — | — | 30272 |
| Dropout (0.3) | (20, 20, 32) | (20, 20, 32) | — | — | 0 |
| Conv2D | (20, 20, 32) | (20, 20, 1) | 3×3 | 1×1 | 290 |
| Total Trainable Parameters: 113345 |  |  |  |  |  |

**Supplementary Table 4:** Discriminator Architecture

| Layer | Input Shape | Output Shape | Filter Size | Stride | #Params |
| --- | --- | --- | --- | --- | --- |
| Input | (2) | (2) | – | – | 0 |
| Dense | (2) | (512) | – | – | 1536 |
| BatchNormalization | (512) | (512) | – | – | 1536 |
| LeakyReLU (0.2) | (512) | (512) | – | – | 0 |
| Dense | (512) | (512) | – | – | 262656 |
| LeakyReLU (0.2) | (512) | (512) | – | – | 0 |
| Dense | (512) | (512) | – | – | 262656 |
| Dropout (0.5) | (512) | (512) | – | – | 0 |
| LeakyReLU (0.2) | (512) | (512) | – | – | 0 |
| Dense | (512) | (1) | – | – | 513 |
| Total Trainable Parameters: 527873 |  |  |  |  |  |

**Supplementary Table 5:**
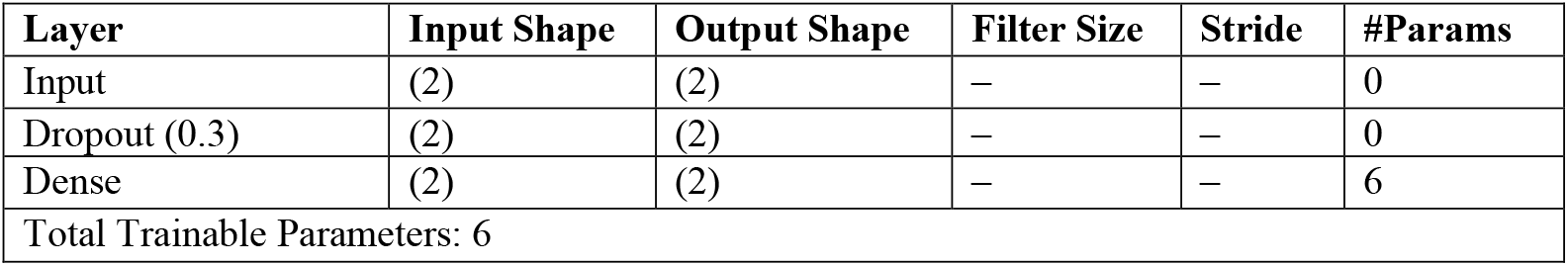
Classifier Architecture

**Supplementary Table 6:** Optimised Classifier for Tabular Features Architecture

| Layer | Input Shape | Output Shape | Filter Size | Stride | #Params |
| --- | --- | --- | --- | --- | --- |
| Input | (2) | (2) | – | – | 0 |
| BatchNormalization | (2) | (2) | – | – | 8 |
| Dense | (2) | (32) | – | – | 96 |
| Dropout (0.3) | (32) | (32) | – | – | 0 |
| Dense | (32) | (16) | – | – | 528 |
| Dropout (0.3) | (16) | (16) | – | – | 0 |
| Dense | (16) | (8) | – | – | 136 |
| Dropout (0.3) | (8) | (8) | – | – | 0 |
| Dense | (8) | (2) | – | – | 18 |
| Total Trainable Parameters: 782 |  |  |  |  |  |

**Supplementary Table 7:** Hand-picked Feature Combination

|  |  |
| --- | --- |
| 1: [FRET_avg, CFP_avg] | 41: [CFP_std, Area] |
| 2: [FRET_avg, YFP_avg] | 42: [CFP_std, Aspect_Ratio] |
| 3: [FRET_avg, CFP_std] | 43: [YFP_std, MajorAxisLength] |
| 4: [FRET_avg, YFP_std] | 44: [YFP_std, MinorAxisLength] |
| 5: [FRET_avg, MajorAxisLength] | 45: [YFP_std, Eccentricity] |
| 6: [FRET_avg, MinorAxisLength] | 46: [YFP_std, Diameter] |
| 7: [FRET_avg, Eccentricity] | 47: [YFP_std, Perimeter] |
| 8: [FRET_avg, Diameter] | 48: [YFP_std, Intensity_avg] |
| 9: [FRET_avg, Perimeter] | 49: [YFP_std, Area] |
| 10: [FRET_avg, Intensity_avg] | 50: [YFP_std, Aspect_Ratio] |
| 11: [FRET_avg, Area] | 51: [MajorAxisLength, MinorAxisLength] |
| 12: [FRET_avg, Aspect_Ratio] | 52: [MajorAxisLength, Eccentricity] |
| 13: [CFP_avg, YFP_avg] | 53: [MajorAxisLength, Diameter] |
| 14: [CFP_avg, CFP_std] | 54: [MajorAxisLength, Perimeter] |
| 15: [CFP_avg, YFP_std] | 55: [MajorAxisLength, Intensity_avg] |
| 16: [CFP_avg, MajorAxisLength] | 56: [MajorAxisLength, Area] |
| 17: [CFP_avg, MinorAxisLength] | 57: [MajorAxisLength, Aspect_Ratio] |
| 18: [CFP_avg, Eccentricity] | 58: [MinorAxisLength, Eccentricity] |
| 19: [CFP_avg, Diameter] | 59: [MinorAxisLength, Diameter] |
| 20: [CFP_avg, Perimeter] | 60: [MinorAxisLength, Perimeter] |
| 21: [CFP_avg, Intensity_avg] | 61: [MinorAxisLength, Intensity_avg] |
| 22: [CFP_avg, Area] | 62: [MinorAxisLength, Area] |
| 23: [CFP_avg, Aspect_Ratio] | 63: [MinorAxisLength, Aspect_Ratio] |
| 24: [YFP_avg, CFP_std] | 64: [Eccentricity, Diameter] |
| 25: [YFP_avg, YFP_std] | 65: [Eccentricity, Perimeter] |
| 26: [YFP_avg, MajorAxisLength] | 66: [Eccentricity, Intensity_avg] |
| 27: [YFP_avg, MinorAxisLength] | 67: [Eccentricity, Area] |
| 28: [YFP_avg, Eccentricity] | 68: [Eccentricity, Aspect_Ratio] |
| 29: [YFP_avg, Diameter] | 69: [Diameter, Perimeter] |
| 30: [YFP_avg, Perimeter] | 70: [Diameter, Intensity_avg] |
| 31: [YFP_avg, Intensity_avg] | 71: [Diameter, Area] |
| 32: [YFP_avg, Area] | 72: [Diameter, Aspect_Ratio] |
| 33: [YFP_avg, Aspect_Ratio] | 73: [Perimeter, Intensity_avg] |
| 34: [CFP_std, YFP_std] | 74: [Perimeter, Area] |
| 35: [CFP_std, MajorAxisLength] | 75: [Perimeter, Aspect_Ratio] |
| 36: [CFP_std, MinorAxisLength] | 76: [Intensity_avg, Area] |
| 37: [CFP_std, Eccentricity] | 77: [Intensity_avg, Aspect_Ratio] |
| 38: [CFP_std, Diameter] | 78: [Area, Aspect_Ratio] |
| 39: [CFP_std, Perimeter] |  |
| 40: [CFP_std, Intensity_avg] |  |

## Supplementary Notes

## Supplementary Note 1: Lung cancer dataset

### Generation of drug-tolerant persister cells for cell fate imaging

On day 1, 5.10^4^ asynchroneous EGFR-mutant non-small cell lung cancer cells (PC9) expressing a nuclear ERK biosensor (EKAREV-NLS) were plated in wells of coverslip-bottomed 24-well plates (ibidi) with 2mL RPMI medium supplemented with 10% Fetal Bovine Serum (FBS). The next day, 500nM EGFR inhibitor (osimertinib) was added to the media. Cells were gently rinsed with PBS and media replaced with fresh osimertinib-containing RPMI twice a week for 2 weeks. While the majority of cells had died, the surviving population consisted of drug-tolerant persister cells (DTPs).

### Time lapse imaging of DTPs

The last media change was performed on the cells approximately 4h prior to imaging. For each biological replicate, between 5 and 9 fields of view were selected. Cells were imaged within an environmental control chamber (5% CO_2_, 37°C) on an Olympus FV3000 laser-scanning confocal microscope with a Plan-Apochromat 20x-0.75NA objective, 445nm laser excitation and 460-500nm and 530-630nm emission windows for CFP and YFP, respectively and a resolution of 1.24µm/pixel. Images were acquired every 4 minutes for 3 days (1080 time points).

### Image processing

Cellular nuclei were segmented using a pre-trained model in ilastik and run on all images using its Fiji plugin. CFP and YFP average nuclear intensities and FRET ratio (YFP/CFP) were computed for each nucleus at each time point, as well as other intensity based (standard deviation of intensities) and morphological features (nuclear area, aspect ratio, etc.) were computed using the ‘regionprops’ function in Matlab. Cells were tracked using Trackmate, allowing for cellular divisions to be detected. Each track was then manually verified for accuracy (tracks with errors were discarded), and cell fate annotation (mitosis or not, time of mitosis).

### Supplementary Note 2: Latent Space Dimension Study

Supplementary Figure 1 examines how the number of latent space dimensions affects model performance. We tested latent dimensions of 2, 3, 5, and 10, measuring model precision across these settings (panel a). While precision remained relatively stable, the highest mean precision was observed when using just two latent features. Moreover, increasing the number of latent dimensions led to higher feature correlations (panel b). With two features, the correlation was minimal (0.06) but adding a third feature slightly increased correlations to 0.10 and 0.05 between feature pairs. As more features were introduced, these correlations became much stronger, reaching 0.68 for five features and exceeding 0.8 for ten features. This suggests that additional features start to capture redundant or less meaningful variations in the data.

A closer examination of the three-feature case revealed that only two features were closely linked to fate classification, while the third feature had little correlation with the task (panel c). Perturbation analysis of this less-relevant feature showed that it primarily captured cell orientation—an attribute that the model learned but that did not contribute meaningfully to classification. In fact, including this feature negatively impacted performance, likely introducing noise rather than useful information. Based on these findings, we selected a two-dimensional latent space, as it effectively captured the two essential and uncorrelated features needed for fate prediction. However, it is important to note that the optimal number of latent features is highly dependent on the specific application. While only two features were critical for classification in this dataset, other biological datasets may require more. This highlights the importance of conducting such analyses to identify which features are both interpretable and relevant for classification.

### Supplementary Note 3: Hand-picked Tabular feature performance

Supplementary Figure 3 compares the model performance of hand-picked features to the latent features from AI4CellFate at the earliest experimental time point. Predictive performance varies widely depending on the chosen features (45%–68% in accuracy) and never outperforms our approach. Panel (a) shows that when using the same classifier as AI4CellFate (with accuracy of 72%), no feature combination exceeds 65% accuracy. Notably, a feature set that performs well with a simple model can perform poorly with a more complex one, highlighting how model choice affects feature importance. Panel (b) further demonstrates this instability: the top features for accuracy differ from those for precision. However, the highest-performing features consistently relate to cell size (diameter, axis length, perimeter) and/or FRET signal—the same features AI4CellFate identified in a fully data-driven manner. Finally, panel (c) shows that even when using all tabular features with an optimised model, accuracy and precision remain below those achieved with AI4CellFate’s latent features, despite the latter using a simpler classifier.

### Supplementary Note 4: Cell Division Times

In this study, we predict whether cells will divide at any point during the 3-day live-cell imaging period. However, it is important to acknowledge that some cells may divide beyond the duration of data acquisition. Supplementary Figure 5 shows the time points at which division occurred, considering only cells labelled as fate 1 (dividing cells). We observe that many divisions happen early, within minutes of acquisition, while fewer occur as time progresses. Even at later time points (e.g., 1000 frames, corresponding to 3 days), a small number of cells (around 7) are still dividing. This suggests that a small percentage of cells labelled as fate 0 (non-dividing) may have been fate 1 but were not captured due to the acquisition window. Consequently, some false predictions of fate 0 could stem from cells that would have eventually divided, but their division occurred outside the recorded timeframe.

## Supplementary Figures

**Supplementary Figure 1.**
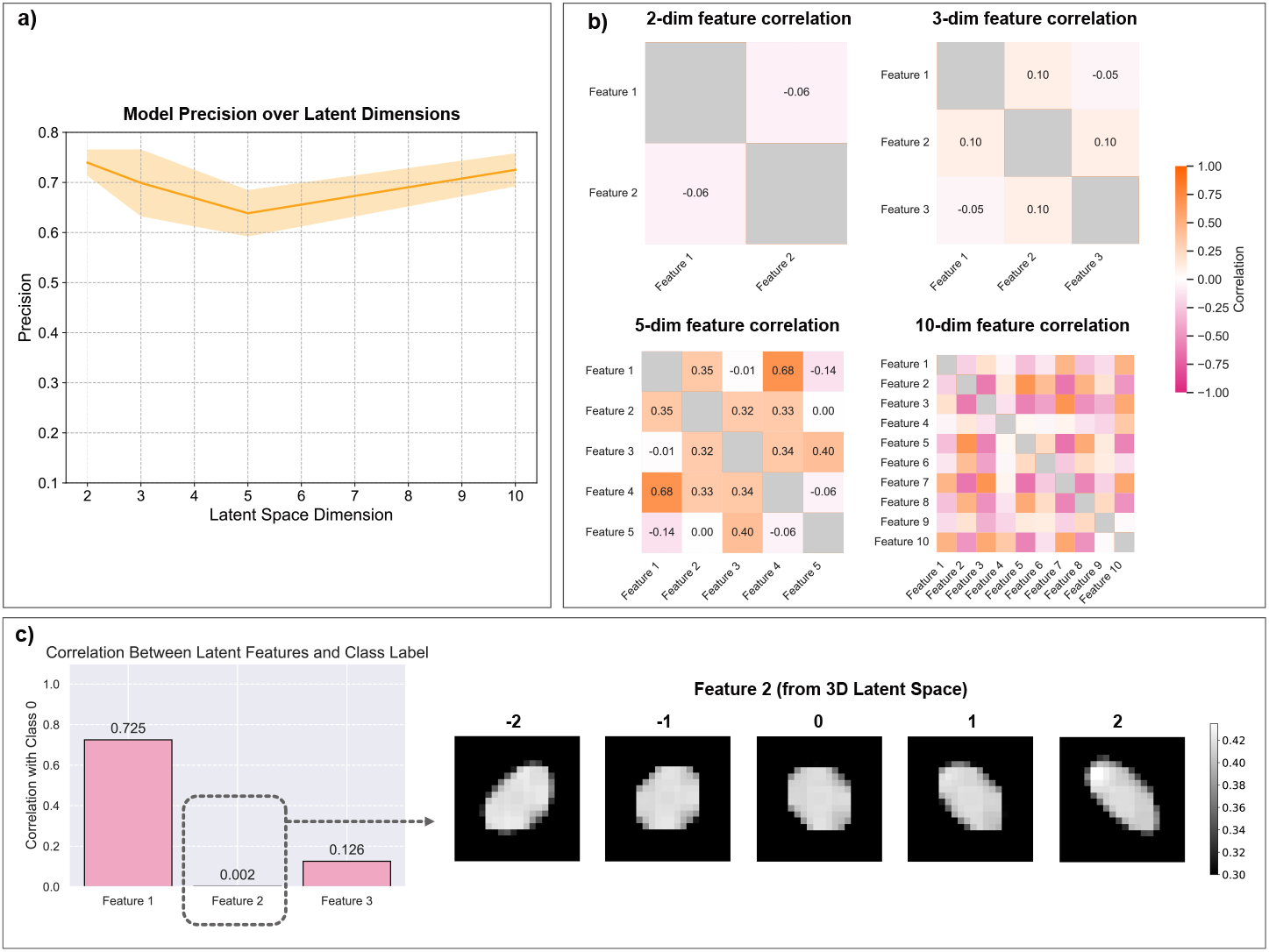
Latent Space Dimensions study. a) Model precision measured for latent space dimensions of 2, 3, 5, and 10 (seeds 42, 43, 44). b) Feature correlation values for the before mentioned dimensions. c) Correlation between latent features and classification task (i.e., fate prediction) on the 3-dimensional latent space case (left). Highlighted is feature 2, showing a low correlation with classification (0.002). (right) Shows the visual perturbations analysis on feature 2.

**Supplementary Figure 2.**
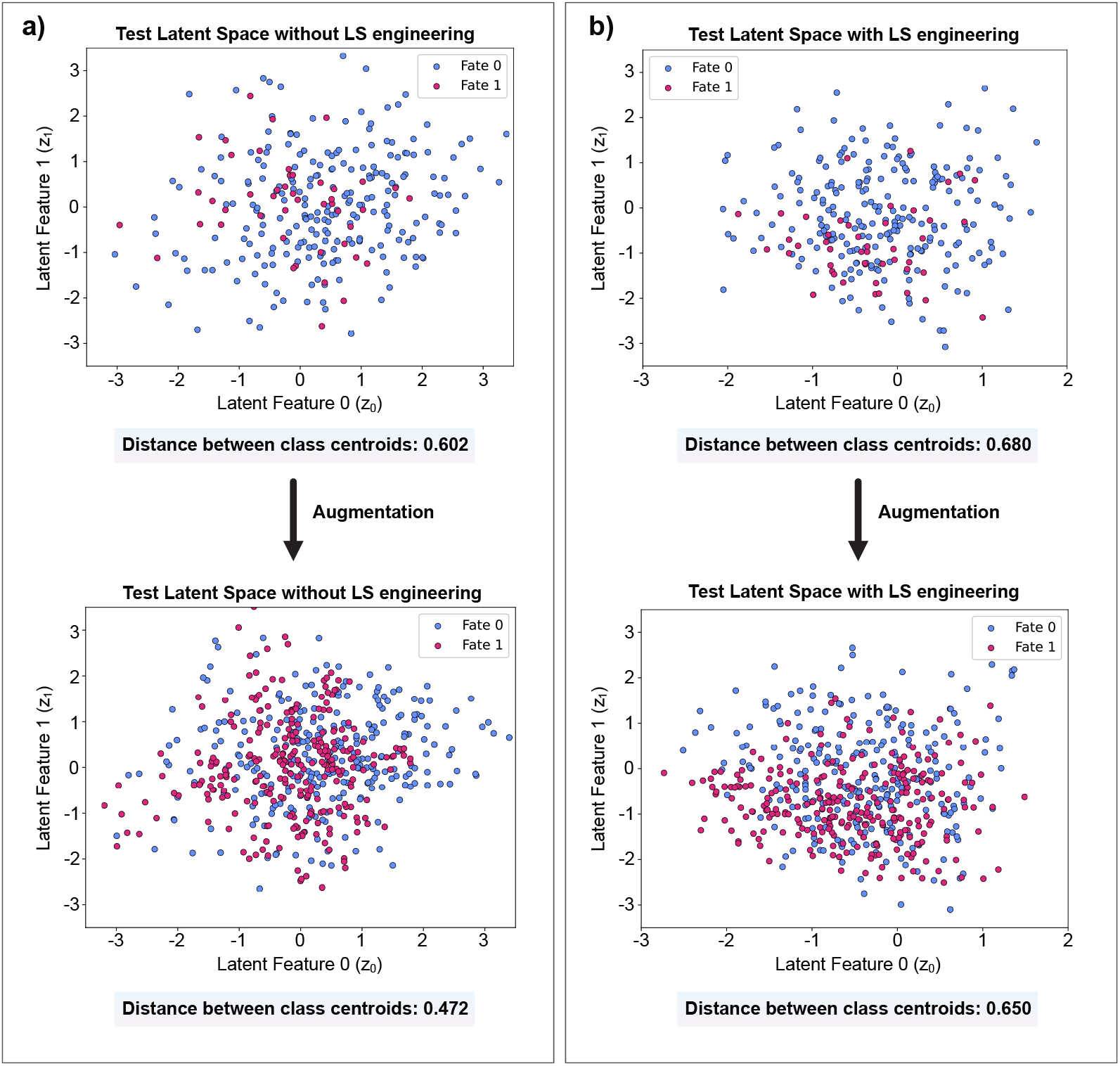
Test Latent Spaces. On top of both panels are the non-augmented test data (original imbalanced test set, 277 cells), and on the bottom are the latent spaces of the augmented test sets (augmented for visualisation purposes only, with the same augmentation function as the training set, described in Methods). a) Test latent spaces without latent space engineering (i.e., without the additional contrastive and covariance losses), showing a distance between class centroids of 0.602 without augmentation and 0.472 with augmentation. b) Test latent spaces with latent space engineering showing a distance between class centroids of 0.680 without augmentation and 0.650 with augmentation.

**Supplementary Figure 3.**
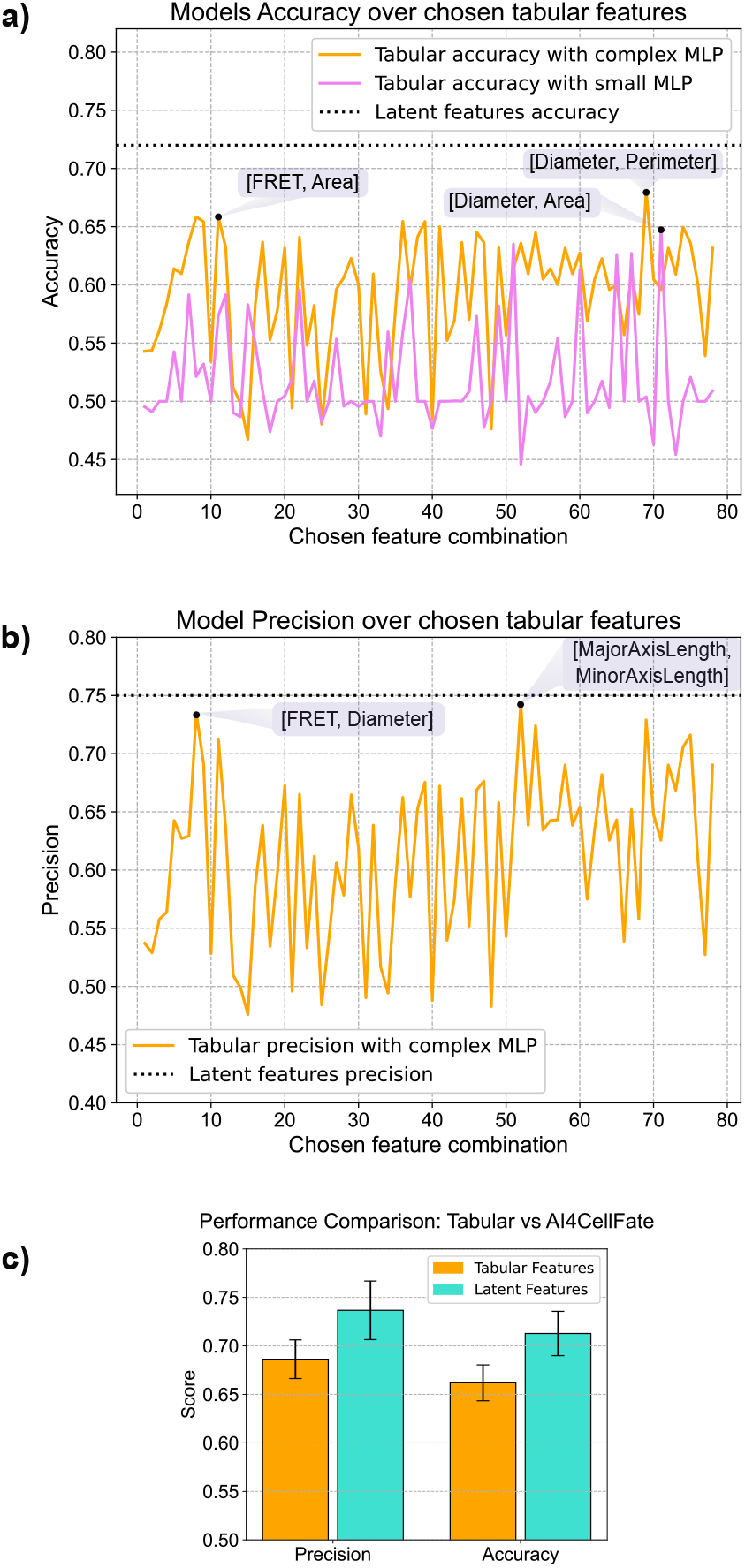
Model performance using hand-picked tabular features in the earliest experimental time point. **(a)** Model accuracy using different hand-picked tabular features, compared to AI4CellFate’s latent features. The orange plot shows accuracy with an optimised MLP, while the pink plot represents a simpler MLP, matching the one used for AI4CellFate. The black dashed line indicates accuracy with AI4CellFate’s features. Highlighted feature sets yielded the highest accuracy. **(b)** Model precision for the optimised MLP on tabular data (orange), with top-performing features highlighted. The black dashed line shows precision with AI4CellFate’s features. **(c)** Comparison of model performance using all tabular features versus AI4CellFate’s latent features. A simple classifier was used for latent features, while the optimised complex classifier was used for tabular features. The left panel shows precision, and the right panel shows accuracy. All results used seed = 42.

**Supplementary Figure 4.**
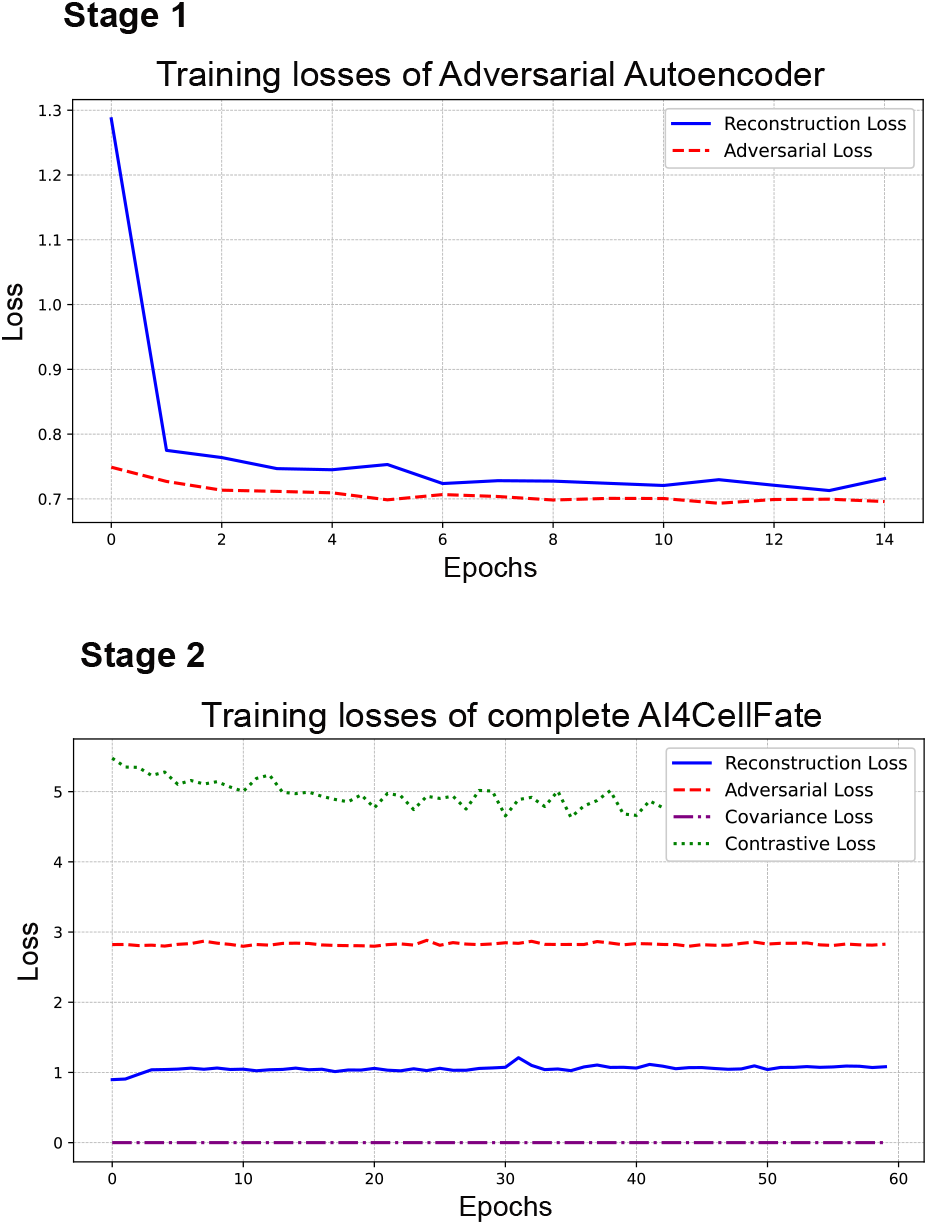
Training loss plots. The first plot (top) includes the training of the adversarial autoencoder only (stage 1), where only the reconstruction and adversarial loss are included (blue and red, respectively). The second plot (bottom) shows the training losses of all four losses used in AI4CellFate (stage 2) – reconstruction, adversarial, covariance and contrastive losses (blue, red, purple, green, respectively).

**Supplementary Figure 5.**
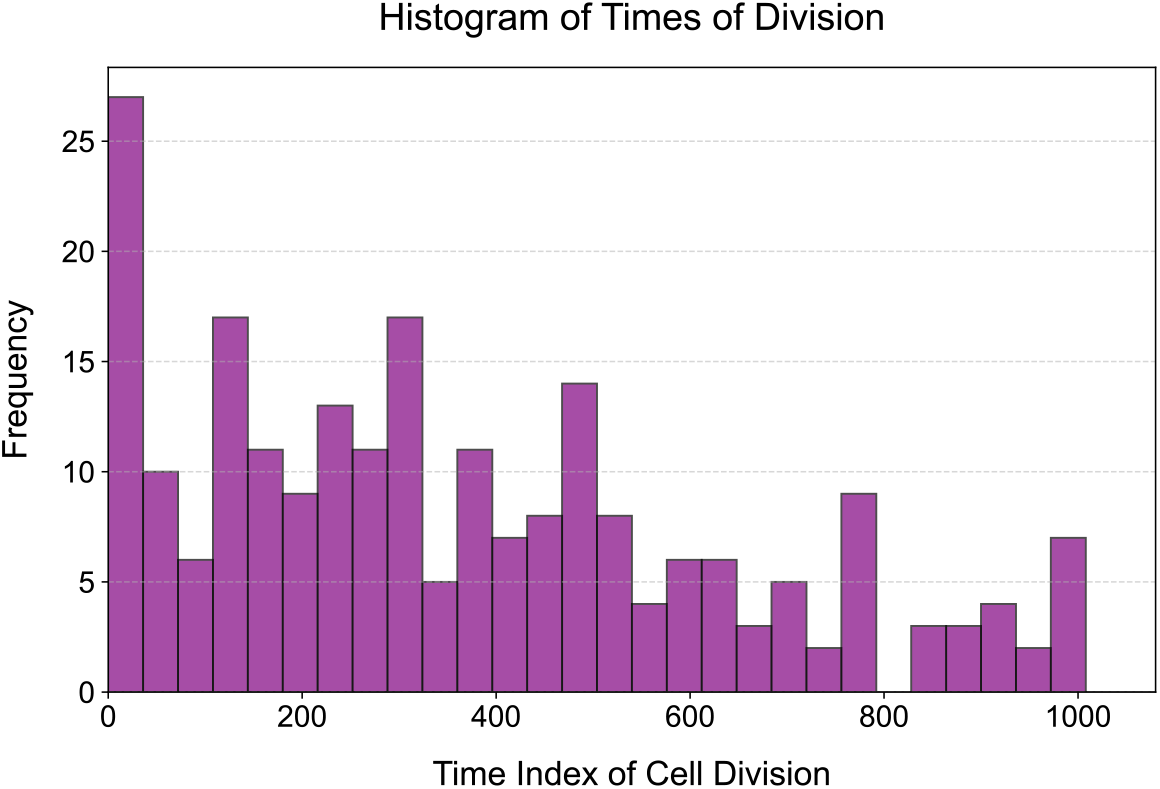
Histogram of time point at which cells divided. The cells used for this histogram were all the cells with fate 1, i.e., the cells that divide at some point in the movie.

